# IL-27 enhances the lymphocyte mediated innate resistance to primary hookworm infection in the lungs

**DOI:** 10.1101/2020.08.12.248021

**Authors:** Jason B. Noon, Arjun Sharma, Johannes Platten, Lee J. Quinton, Christoph Reinhardt, Markus Bosmann

## Abstract

Interleukin-27 (IL-27) is a heterodimeric cytokine of the IL-12 family, formed by non-covalent association of the promiscuous EBI3 subunit and selective p28 subunit. IL-27 is produced by mononuclear phagocytes and unfolds pleiotropic immune-modulatory functions through high affinity ligation to IL-27 receptor alpha (IL-27RA). While IL-27 is known to contribute to immunity and to end inflammation following numerous types of infections, its relevance for host defense against multicellular parasites is still poorly defined. Here, we investigated the role of IL-27 during infection with the soil-transmitted hookworm, *Nippostrongylus brasiliensis,* in its early intrapulmonary life cycle. IL-27(p28) was detectable in broncho-alveolar lavage fluids of C57BL/6J wild type mice on day 1 after subcutaneous *N. brasiliensis* inoculation. The expression of IL-27RA was most abundant on lung invading γδ T cells followed by CD8^+^ T cells, CD4^+^ T cells and NK cells. IL-27RA was weakly present on CD19^+^ B cells and absent on neutrophils, alveolar macrophages and eosinophils. Il27ra^−/−^ mice showed increased parasite burden together with aggravated pulmonary hemorrhage and higher alveolar albumin leakage as a surrogate for disruption of the epithelial/vascular barrier. Conversely, recombinant mouse IL-27 injections of wild type mice reduced parasite burdens and lung injury. In multiplex screens, we identified higher airway accumulations of IL-6, TNFα and MCP-3 (CCL7) in Il27ra^−/−^ mice, while rmIL-27 treatment showed a reciprocal effect. Finally, γδ T cell infiltration of the airways required endogenous IL-27 expression. In summary, this report demonstrates protective functions of IL-27 to control the early larval stage of hookworm infection in the lungs.

## Introduction

Hookworms are soil-transmitted intestinal nematodes that have a critical stage in development within the airspaces of the lungs (1). After molting from an infectious third stage (L3) to an L4 stage within the alveolar space, hookworms ascend the trachea and are swallowed, ultimately infecting the small intestine where they remain as egg-laying adults. In the small intestine, hookworms rupture vessels and feed on blood, which is the cause of clinical hookworm disease characterized by iron-deficiency anemia. Over 500 million people worldwide are infected with hookworms (2, 3), and among all parasites, hookworms are behind only malaria for the leading causes of iron-deficiency anemia globally (4–7), indicating the importance of these parasites. Although there are anthelmintic drugs available for treating hookworms (8, 9), people are rapidly re-infected in endemic areas due to insufficient immunity and high vulnerability for secondary infections (10). Importantly, there is not a single licensed hookworm vaccine (11), stressing the value of understanding protective immunity to hookworm infections.

Murine hookworm *Nippostrongylus brasiliensis* is a model for human hookworm disease, particularly for the stage of infection within the lungs, which occurs on days 1 and 2 after subcutaneous inoculation (12). On day 3, larvae transition to the small intestine and remain until day 7. There are many publications that indicate the importance of the canonical type 2 response involving cytokines such as IL-4, IL-5, IL-13 and RELMβ, along with group 2 innate lymphoid cells (ILC2s), type 2 T helper (Th2) cells, alternatively activated macrophages, eosinophils, basophils, mast cells and goblet cells in resistance to secondary *N. brasiliensis* infections in the lungs (13), which does not occur in humans. However, knowledge on primary *N. brasiliensis* infection in the lungs is sparse (14, 15). Resistance to primary hookworm infections (in animals) is widely believed to be localized to the small intestine stage of infection and, in the *N. brasiliensis* model, is well-characterized by canonical type 2-driven expulsion mediated by ILC2s and Th2 cells (13), as well as type 2 γδ T (γδ T2) cells (i.e., intestinal intraepithelial T lymphocytes; IELs) (16). Although much less is known about primary resistance in the lungs to hookworm infections, in the *N. brasiliensis* model, IL-17A, neutrophils and γδ T cells (14), and interestingly, ILC2s (15) are involved.

IL-27 is a heterodimeric cytokine composed of a unique p28 α-subunit and an EBI3 β-subunit (17). EBI3 is shared with IL-35 (18). The IL-27 receptor is also a heterodimer composed of a unique IL-27 receptor α-subunit (IL-27RA, WSX-1) and a gp130 β-subunit that is shared with multiple other cytokine receptors (17, 19, 20). IL-27 is well-described to exert acute pro-inflammatory effects, enhancing type 1 responses, particularly CD8^+^ cytotoxic lymphocytes (CTLs) and natural killer (NK) cells, thus enhancing protective immunity to intracellular pathogens and various cancers (21). Consistent with IL-27 enhancing type 1 responses, IL-27 directly suppresses the expansion and activation of ILC2s during the lung repair phase of primary *N. brasiliensis* infections (22). A more rapid intestinal expulsion is seen in the absence of IL-27 activities in both models of hookworm (studying Ebi3^−/−^ mice) and whipworm (studying Il27ra^−/−^ mice) infections (22, 23). Importantly, there are no publications on IL-27 during early primary *N. brasiliensis* infections in the lungs, or any other helminth infection. As ILC2s are involved in limiting primary *N. brasiliensis* infections in the lungs (24, 25) and IL-27 antagonizes tissue-resident ILC2s (26), we initially hypothesized that IL-27 limits resistance to primary *N. brasiliensis* infections in the lungs by suppressing type 2 responses. Interestingly, however, we found that IL-27 enhances resistance to primary *N. brasiliensis* infections in the lung alveolar space in association with increased expansion of γδ T cells, which we also found to express much higher levels of IL-27RA compared both CD4^+^ Th cells and CD8^+^ CTLs.

In conclusion, future efforts for the development of a hookworm vaccine may be inspired by the concept of targeting the lung larval stage and considering adjuvants likely to induce strong IL-27-dependent immunity (11, 27–29).

## Materials and Methods

### Mice

All procedures with mice were approved by the Institutional Animal Care and Use Committee of the Boston University and performed in compliance with the guidelines of the National Institutes of Health. IL-27RA^−/−^ mice (B6N.129P2-Il27ra^tm1Mak^/J; on C57BL/6NJ background), C57BL/6NJ mice and C57BL/6J mice were obtained from the Jackson Laboratory (Bar Harbor, ME). The mice were bred and genotyped at the animal facilities of the Boston University under specific pathogen-free conditions and controlled light/dark cycle. Male and female mice at 8-12 weeks of age were used for experiments.

### Nippostrongylus brasiliensis cultures and inoculations

*N. brasiliensis* coprocultures were provided by Dr. Joseph Urban Jr., United States Department of Agriculture. L3 were extracted from coprocultures and prepared for inoculations according to an established protocol (12). Mice were injected subcutaneously in the flank with 500 L3 in 0.1 mL of sterile PBS (Thermo Fisher Scientific, Waltham, MA).

### Alveolar and lung tissue parasite burdens

To measure alveolar parasite burden, inoculated mice were euthanized by CO_2_ overdose, and three bronchoalveolar lavages (BAL) were collected before vital organ removal. For the first BAL, 1 mL PBS containing 1X HALT Protease Inhibitor Cocktail/EDTA (Thermo Fisher Scientific) was collected into a 1.5 mL tube. PBS only was used for the second and third BAL and were collected into a 50 mL tube on ice. The first BAL was centrifuged at 400 x g for 8 min at room temperature and the cell-free supernatant was collected as BAL fluid (BALF) and stored at −80°C for later analysis. The pellet of BAL cells and containing *N. brasiliensis* larvae was then resuspended in PBS and transferred to the 50 mL tube along with the additional BAL collections. The combined BAL was then diluted to 30 mL with PBS and transferred to a 100 mm petri dish with grids drawn on the bottom surface. All intact *N. brasiliensis* larvae were counted under 20X magnification (AmScope, Irvine, CA), excluding *N. brasiliensis* debris and obviously dead and deteriorating larvae. The suspension of BAL cells and *N. brasiliensis* larvae was then transferred back to the 50 mL tube and centrifuged at 800 x g for 8 min at 4°C. The supernatant was removed down to 10 mL, then the BAL cell pellet was resuspended, transferred to a 15 mL tube and centrifuged as before. The supernatant was removed down to 0.5 mL, then the BAL cell pellet was transferred to a 1.5 mL tube before further analysis. To measure parasite burden in lung tissues, lungs were collected from inoculated mice after BAL collections into 35 mm petri dishes. Lungs were minced with a surgical scissors, resuspended in 5 mL of a 37°C slurry of 1% agarose, and then pipetted onto a flattened layer of cheesecloth. Once solidified, the cheesecloth was rolled up and placed in a submerging vessel of 45 mL PBS in a 50 mL tube, with a small piece of the cheesecloth secured between the tube and lid, and then incubated in 37°C overnight. Larvae that migrated out of the lung tissues and agarose were counted under 20X magnification.

### Alveolar injury

BALF total protein was measured with a Pierce BCA Protein Assay Kit (Thermo Scientific). Alveolar hemorrhage was assessed as described elsewhere (30), but with the following modifications. A 6X 2-fold serial dilution (8000-250 μg/mL) of human hemoglobin (Sigma, St. Louis, MO) was prepared, and 50 μL of each standard and BAL sample was added to duplicate wells of a 96-well plate. A volume of 100 μL of 6% sodium dodecyl sulfate (Sigma) was added to all wells and resuspended several times. The absorbance at 560 nm was measured with a Tecan Infinite M Nano plate reader (Tecan, Männedorf, Switzerland), and alveolar hemorrhage (i.e., total amount of hemoglobin recovered in BAL) was determined from the standard curve generated by Magellan V 7.2 (Tecan) software.

### ELISA & multiplex bead-based immunoassay

IL-27(p28) in BALF was measured with a mouse IL-27 p28/IL-30 DuoSet ELISA (R&D Systems, Minneapolis, MN), according to the manufacturer’s instructions. The absorbance at 450 nm was measured with a Tecan Infinite M Nano plate reader, and concentration was determined from the standard curve generated by Magellan software.

A multiplex bead-based immunoassay (Cytokine & Chemokine 26-Plex Mouse ProcartaPlex™ Panel 1, Thermo Fisher Scientific) was used for simultaneous quantification of the following cytokines/chemokines: IL-1β, IL-2, IL-4, IL-5, IL-6, IL-9, IL-10, IL-12p70, IL-13, IL-17A, IL-18, IL-22, IL-23, IL-27, GROα (CXCL-1), IP-10 (CXCL-10), MCP-1 (CCL-2), MCP-3 (CCL-7), MIP-1α (CCL-3), MIP-1β (CCL-4), MIP-2 (CXCL-2), RANTES (CCL-5), Eotaxin (CCL-11), GM-CSF, IFN gamma, and TNF alpha (31). All samples from bead-based assays were performed using a LiquiChip-200 instrument (Qiagen, Hilden, Germany) using Bio-Plex Manager v6.1 software for quantification.

### Flow Cytometry

Fresh BAL cells were collected from inoculated mice as described above. In multiple experiments using our standard operating procedures, >99% of BAL cells were found to be negative for fixable viability dye eFlour 780 (eBioscience, ThermoFisher Scientific), and hence, all BAL cells are considered alive. BAL cells collected in a 15 mL tube were centrifuged 800 x g at 4°C for 8 min, and then resuspended in the following mouse antibody staining cocktails, transferred to 1.5 mL tubes, and then incubated in the dark for 30 min at 4°C: Myeloid-lymphocyte common lineage panel – TruStain FcX (anti-CD16/32) (clone: 93; dilution: 1:100, BioLegend, San Diego, CA), CD45-Pacific Blue (clone: 30-F11; dilution 1:200, BioLegend), Ly6G-APC (clone: 1A8; dilution: 1:400, BioLegend), Siglec-F-APC/Cy7 (clone: E50-2440; dilution: 1:200, BioLegend), CD11c-Alexa Fluor 488 (clone: N418; dilution: 1:600, BioLegend), CD3-BUV737 (clone: 145-2C11; dilution: 1:100, BD Biosciences, San Jose, CA), CD19-PE/Cy7 (clone: 6D5; dilution: 1:100, BioLegend), NK1.1-PerCP/Cy5.5 (clone: PK136; dilution: 1:100, BioLegend), IL-27RA-PE (clone: 2918; dilution: 1:100, BD Biosciences); T cell panel – TruStain FcX (anti-CD16/32) (clone: 93; dilution: 1:100, BioLegend), CD3-BUV737 (clone: 145-2C11; dilution: 1:100, BD Biosciences), TCRβ chain-PE/Cy7 (clone: H57-597; dilution: 1:100, BioLegend), CD4-Alexa Fluor 488 (clone: RM4-5; dilution 1:400, BioLegend), CD8a-APC/Cy7 (clone: 53-6.7; dilution: 1:100, BioLegend), TCRγδ-APC (clone: GL3; dilution: 1:00, BioLegend), NK1.1-PerCP/Cy5.5 (clone: PK136; dilution: 1:100, BioLegend), IL-27RA-PE (clone: 2918; dilution: 1:100, BD Biosciences) or PE Rat IgG2a, κ Isotype Control (clone: R35-95; dilution: 1:100, BD Biosciences). All antibodies were diluted in FACS buffer prepared from sterile PBS and supplemented with 0.25% (w/v) BSA, 0.02% (w/v) sodium azide and 2 mM EDTA. The stained cells were rinsed in

FACS buffer, and then fixed in 2% paraformaldehyde (Santa Cruz Biotechnology, Dallas, TX) at room temperature for 20 min. Stained/fixed cells were centrifuged as described before, resuspended in FACS buffer and transferred to a FACS tube containing 20 μL of CountBright Absolute Counting Beads (ThermoFisher Scientific). For all single-stained compensation controls, we used OneComp eBeads Compensation Beads (Invitrogen) according to the manufacturer’s instructions. Flow cytometric analysis was performed on a BD LSR II flow cytometer with BD FACSDiva software. Final plots were made in FlowJo v10.

### Reagents

Recombinant mouse IL-27 (<1.0 EU per 1 μg of the rmIL-27 protein by the LAL method) was purchased from R&D systems.

### Data analysis

Statistical analyses were performed and graphs were prepared in Prism v7.04-v8.4.3 (GraphPad Software). Data in bar graphs are depicted as mean ± standard error of the mean (S.E.M.), with overlaid symbols representing values from individual mice. Two-group single comparisons were made with a t-test (with parametric data pre-confirmed using the F test). Multiple comparisons were made with a one-way ANOVA followed by Tukey’s post hoc test. *P* values <0.05 were considered significant. *P* values <0.1 were considered as a trend.

## Results

### IL-27(p28) is transiently elevated in alveolar space during primary N. brasiliensis infection of the lungs

While *N. brasiliensis* infection is well known, to induce an eosinophilic and Th2 cell dependent immune response, the role of the immune-modulatory cytokine IL-27 is not well described in the pulmonary state of the hookworm infection cycle. To test if IL-27(p28) is released during primary *N. brasiliensis* infection of the lungs, C57BL/6J mice (wild type, WT) were inoculated with n=500 *N. brasiliensis* L3, and EDTA-plasma was collected on days 0, 1, 2, 6 and 9 following infection. BALF was collected on days 0, 1, 2 and 9 post infection (p.i.). Circulating IL-27(p28) was not detectable in EDTA-plasma by ELISA at any of the studied time points, even though all mice were confirmed to be highly infected with an average of 40,000 eggs per gram of feces on day 6 p.i. (data not shown). However, in BALF, IL-27(p28) was elevated by ~2.6-fold on day 1 p.i. compared to day 0 (*P*<0.0001, Fig. 1). The concentrations of IL-27(p28) returned to normal levels by day 2 (Fig. 1). Thus, these results indicate that IL-27(p28) is transiently released specifically in the alveolar space during primary *N. brasiliensis* infection of the lungs. The bulk of IL-27 may avidly bind to its receptor for rapid clearance or altogether escape detection in cell-free BALF.

**FIGURE 1.**
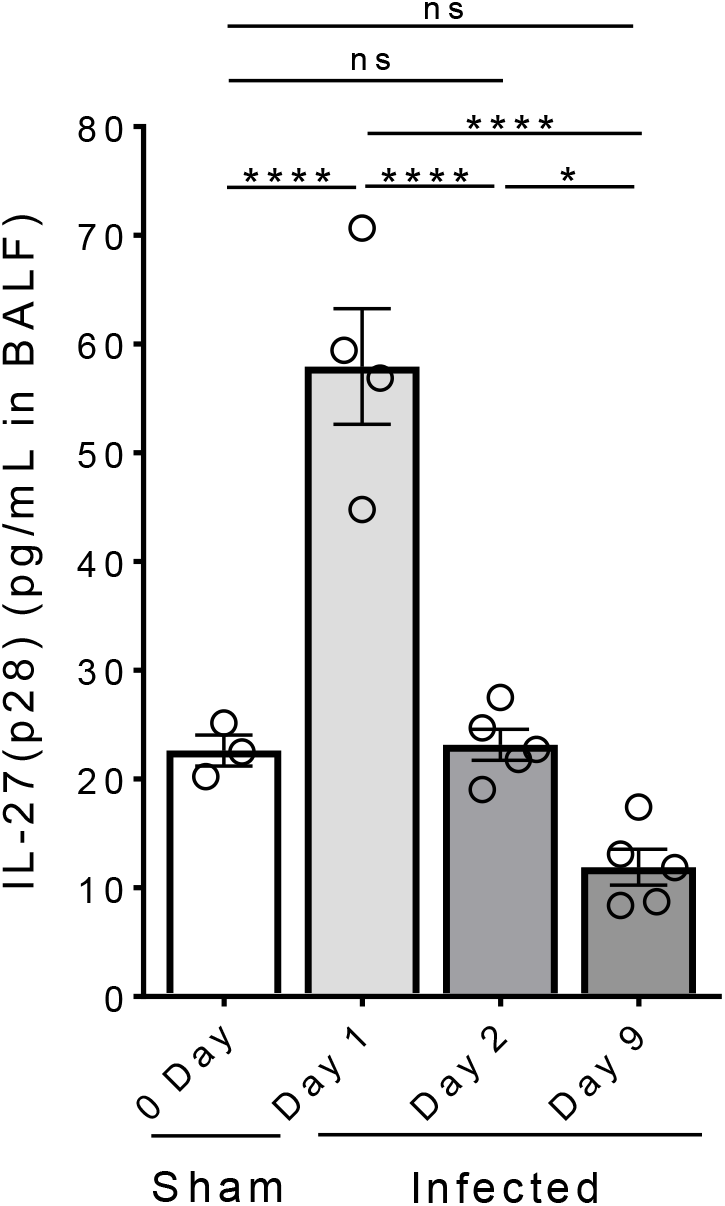
IL-27(p28) is transiently present in the lung alveolar space at day 1 post-inoculation with *N. brasiliensis* larvae. Wild type mice (C57BL/6J) were inoculated s.c. with L3 larvae (n=500/mouse) of *N. brasiliensis.* IL-27(p28) was quantified in broncho-alveolar lavage fluids (BALF) by ELISA at the indicated time points (days 0, 1, 2 and 9). Comparisons of mean ± SEM and each circle represents an individual animal. PBS: mock inoculated/uninfected/day 0 post-inoculation. * *P*<0.05, **** *P*<0.0001, ns: not significant.

### IL-27RA is highly expressed on T cells in alveolar space during primary pulmonary N. brasiliensis infection

To study if IL-27RA is expressed on invading immune cells in the broncho-alveolar space during primary *N. brasiliensis* infection of the lungs, we first confirmed leukocyte dynamics in BAL from C57BL/6J mice on days 0, 1 and 2 p.i. with *N. brasiliensis* larvae. Within the myeloid lineage, we recovered an average of ~70,000 CD11c^+^Siglec-F^+^ resident alveolar macrophages per mouse in BAL on day 0 in sham mice (Fig. S1A, S1B). The numbers of alveolar macrophages were similar in BAL on day 1 p.i., while recovery of this cell type increased to ~120,000 in BAL by day 2 p.i. (Fig. S1A, S1B). In addition, while no Ly6G^+^ neutrophils or CD11c^−^Siglec-F^+^ eosinophils were present in BAL of sham mice, the numbers of these myeloid cells substantially increased over the course of 2 days following *N. brasiliensis* infection (Fig. S1B). Furthermore, within the lymphocyte lineage, while few cells were detectable in day 0 p.i. BAL, accumulation of CD19^+^CD3^−^ B cells, CD19^−^CD3^+^ T cells and CD19^−^CD3^−^NK1.1^+^ NK cells was detected on day 1 and further elevated on day 2 (Fig. 2A, 2B). Thus, all investigated myeloid and lymphoid cells increase in the alveolar space during the natural course of primary *N. brasiliensis* infection.

**FIGURE 2.**
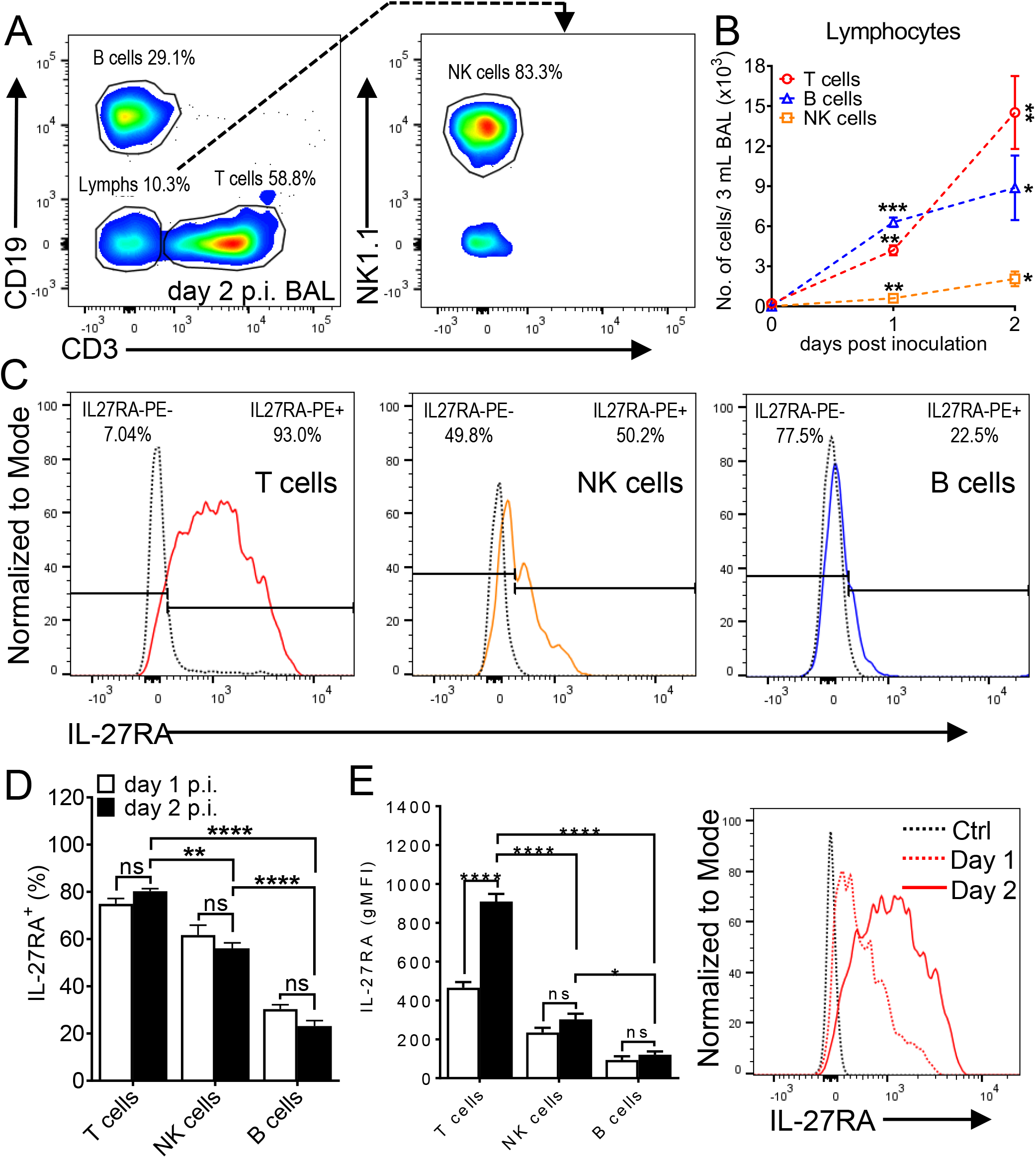
*N. brasiliensis* infection promotes the appearance of IL-27RA expressing lymphocytes in the lungs. C57BL/6J wild type mice were infected s.c. with L3 larvae (n=500/mouse) or received a mock PBS injection as controls. The inflammatory cells were collected by BAL at indicated time points and analyzed by flow cytometry. (**A**) Plots of CD19 versus CD3 pre-gated on CD45^+^Ly6G^−^ single cell lymphocytes (left panel) and NK1.1 versus CD3 gated on CD19^−^CD3^−^ innate lymphocytes (right panel) at 2 days after infection. Percentages are indicated next to each gate. (**B**) Absolute numbers of lymphocyte populations in BAL of CD19^−^CD3^+^ T cells, CD19^+^CD3^−^ B cells and CD19^−^CD3^−^NK1.1^+^ NK cells on day 0 (PBS-inoculated/uninfected), day 1 and day 2 post-inoculation (n=3 mice/group). (**C**) Representative histograms of IL-27RA expression on CD19^−^CD3^+^ T cells. (left panel), CD19^−^CD3^−^NK1.1^+^ NK cells (middle panel) and CD19^+^CD3^−^ B cells (right panel) on day 2 p.i. The dotted black line indicates isotype-FMO control. The percentages of IL-27RA^−^ and IL-27RA^+^ cells are indicated in the upper left and right corners, respectively. (**D**) Frequencies (%) of IL-27RA^+^CD19^−^CD3^+^ T cells, CD19^−^CD3^−^NK1.1^+^ NK cells and CD19^+^CD3^−^ B cells in BAL on day 1 (n=2 mice/group) and day 2 (n=3 mice/group) post-inoculation. (**E**) IL-27RA presence expressed as geometric mean fluorescence intensities (gMFI) on CD19^−^CD3^+^ T cells, CD19^−^CD3^−^NK1.1^+^ NK cells and CD19^+^CD3^−^ B cells (all left panel) from the same experiments described in frame D. A representative histogram of IL-27RA expression on CD19^−^CD3^+^ T cells is shown in the right panel with the dotted and solid red lines indicating days 1 and 2 p.i., respectively. The dotted black line indicates isotype-FMO control (Ctrl). Data (B, D, E) are shown as mean ± SEM and were analyzed by two-tailed t-test (B) comparing day 0 vs. day 1 and day 0 vs. day 2 for each cell type, or two-way ANOVA (D, E), * *P*<0.05, ** *P*<0.01, *** *P*<0.001, **** *P*<0.0001, ns: not significant.

Essentially none of the three myeloid cell types (neutrophils, eosinophils, alveolar macrophages) present in BAL after infection were found to express IL-27RA at any time point evaluated (Fig. S1C). However, an average of 80.2%, 56.1% and 23.1% of CD19^−^CD3^+^ T cells, CD19^−^CD3^−^NK1.1^+^ NK cells and CD19^+^CD3^−^ B cells were IL-27RA^+^, and these percentages were similar on both day 1 and 2 p.i. (Fig. 2C, 2D). Geometric mean fluorescence intensities (gMFI) of IL-27RA were 2-fold greater on CD19^−^CD3^+^ T cells on day 2 compared to day 1 p.i. (suggesting T cell-specific upregulation of IL-27RA, Fig. 2E), while IL-27RA gMFI was similar on CD19^−^ CD3^−^NK1.1^+^ NK cells and CD19^+^CD3^−^ B cells at the two time points studied (Fig. 2E). Moreover, IL-27RA gMFI was >3-fold higher on CD19^−^CD3^+^ T cells compared to CD19^−^CD3^−^NK1.1^+^ NK cells and CD19^+^CD3^−^ B cells. Hence, the frequency and magnitude of IL-27RA expression is greatest on CD19^−^CD3^+^ T cells in the alveolar space during primary *N. brasiliensis* infection of the lungs.

We next evaluated the numbers of different T cell subsets in the alveolar space during primary pulmonary *N. brasiliensis* infection, and for their expression of IL-27RA. Both TCRβ^+^CD4^+^ Th cells and TCRβ^+^CD8^+^ CTLs became considerably more abundant than TCRβ^−^ TCRγδ^+^ γδ T cells after *N. brasiliensis* infection, while none of these lymphocyte populations were present in BAL from uninfected mice (Fig. 3A, 3B). The numbers of TCRβ^+^CD4^+^ Th cells, TCRβ^+^CD8^+^ CTLs and TCRβ^−^TCRγδ^+^ γδ T cells further increased from day 1 to day 2 (Fig. 3A, 3B). Moreover, >40% of all three subsets were IL-27RA^+^ (Fig. 3C, 3D). There was a significantly higher percentage of IL-27RA^+^TCRβ^−^TCRγδ^+^ γδ T cells compared to IL-27RA^+^TCRβ^+^CD8^+^ CTLs and IL-27RA^+^TCRβ^+^CD4^+^ Th cells, while there was not a significant difference between IL-27RA^+^TCRβ^+^CD8^+^ CTLs and IL-27RA^+^TCRβ^+^CD4^+^ Th cells (Fig. 3D). Furthermore, TCRβ^−^ TCRγδ^+^ γδ T cells resulted in significantly higher IL-27RA gMFI compared to both TCRβ^+^CD8^+^ CTLs and TCRβ^+^CD4^+^ Th cells, and TCRβ^+^CD8^+^ CTLs resulted in significantly higher IL-27RA gMFI compared to TCRβ^+^CD4^+^ Th cells (Fig. 3E). Thus, IL-27RA is most prevalent and most highly expressed on γδ T cells, but Th cells and CTLs dominate the T cell population in the alveolar space during primary *N. brasiliensis* infection of the lungs.

**FIGURE 3.**
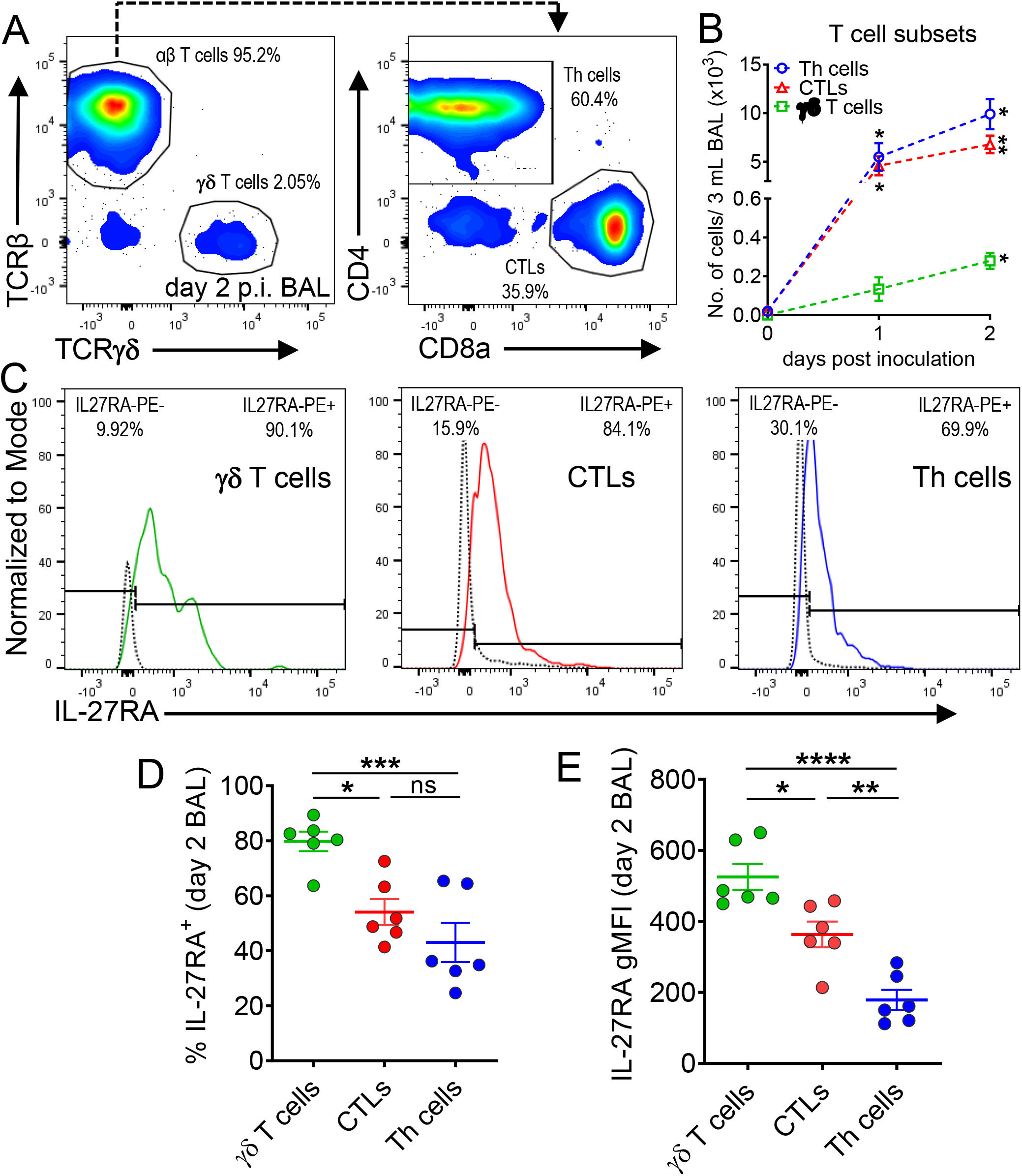
IL-27RA expression is highest on γδ T cells among broncho-alveolar lymphocytes during *N. brasiliensis* infection. (**A**) Flow cytometry plots of TCRβ chain versus TCRγδ pre-gated on CD3^+^ single cell lymphocytes (left panel) and CD4 versus CD8a on αβ T cells (right panel) in BAL after 2 days of infection with L3 larvae of *N. brasiliensis* (n=500 per C57BL/6/J mouse s.c.). The frequencies (%) of cells are indicated next to each gate. (**B**) Absolute numbers in BAL of CD4^+^CD8^−^ T helper (Th) cells, CD4^−^ CD8^+^ cytotoxic T lymphocytes (CTLs) and TCRβ^−^TCRγδ^+^ γδ T cells on day 0 (PBS-inoculated/uninfected), day 1 (n=2 mice/group) and day 2 post-inoculation (n=6 mice/group). (**C**) Representative histograms of IL-27RA expression on TCRβ^−^TCRγδ^+^ γδ T cells (left panel), CD4^−^ CD8^+^ CTLs (middle panel) and CD4^+^CD8^−^ Th cells (right panel). The dotted line indicates IL-27RA staining in Il27ra^−/−^ mice as negative control. The percentages of IL-27RA^−^ and IL-27RA^+^ cells are indicated in the upper left and right corners, respectively. (**D**) Relative numbers of IL-27RA-positive TCRβ^−^TCRγδ^+^ γδ T cells, CD4^−^CD8^+^ CTLs and CD4^+^CD8^−^ Th cells in BAL on day 2 p.i. (n=6 mice/group). (**E**) IL-27RA abundance (gMFI) on TCRβ^−^TCRγδ^+^ γδ T cells, CD4^−^CD8^+^ CTLs and CD4^+^CD8^−^ Th cells in BAL on day 2 p.i. (n=6 mice/group). Data (B, D, E) are shown as mean ± SEM and were analyzed by two-tailed t-test (B) comparing day 0 vs. day 1 and day 0 vs. day 2 for each cell type, or one-way ANOVA (D, E), * *P*<0.05, ** *P*<0.01, *** *P*<0.001, **** *P*<0.0001, ns: not significant.

### IL-27 signaling enhances resistance to primary N. brasiliensis infections in the lungs

To determine if IL-27 has a beneficial or detrimental role during primary *N. brasiliensis* infection of the lungs, we first inoculated Il27ra^−/−^ mice and C57BL/6NJ wild type (WT) mice with 500 *N. brasiliensis* L3 and compared alveolar parasite burden, hemorrhage and total protein on day 2 p.i.. Strikingly, Il27ra^−/−^ mice showed a 2.2-fold increase in alveolar parasite burdens compared to WT mice (Fig. 4A). We confirmed that the majority of parasites are recovered by the BAL procedure with only few remaining larvae in lung tissues (Fig. S2). To evaluate the severity of parasite-induced lung injury, we determined hemoglobin concentrations in BAL as a marker for airway hemorrhage and broncho-alveolar albumin as a surrogate endpoint for the disturbance of the epithelial/vascular barrier function. A significant 1.9-fold increase in alveolar hemorrhage in Il27ra^−/−^ mice compared to WT mice was observed in line with the greater parasite burden of Il27ra^−/−^ mice (Fig. 4B). Moreover, and also consistent with the greater parasite burden, a moderate but significant 1.3-fold increase in total protein leakage was detected in the BALF of Il27ra^−/−^ mice (Fig. 4C), indicating an increase in proteinaceous edema.

**FIGURE 4.**
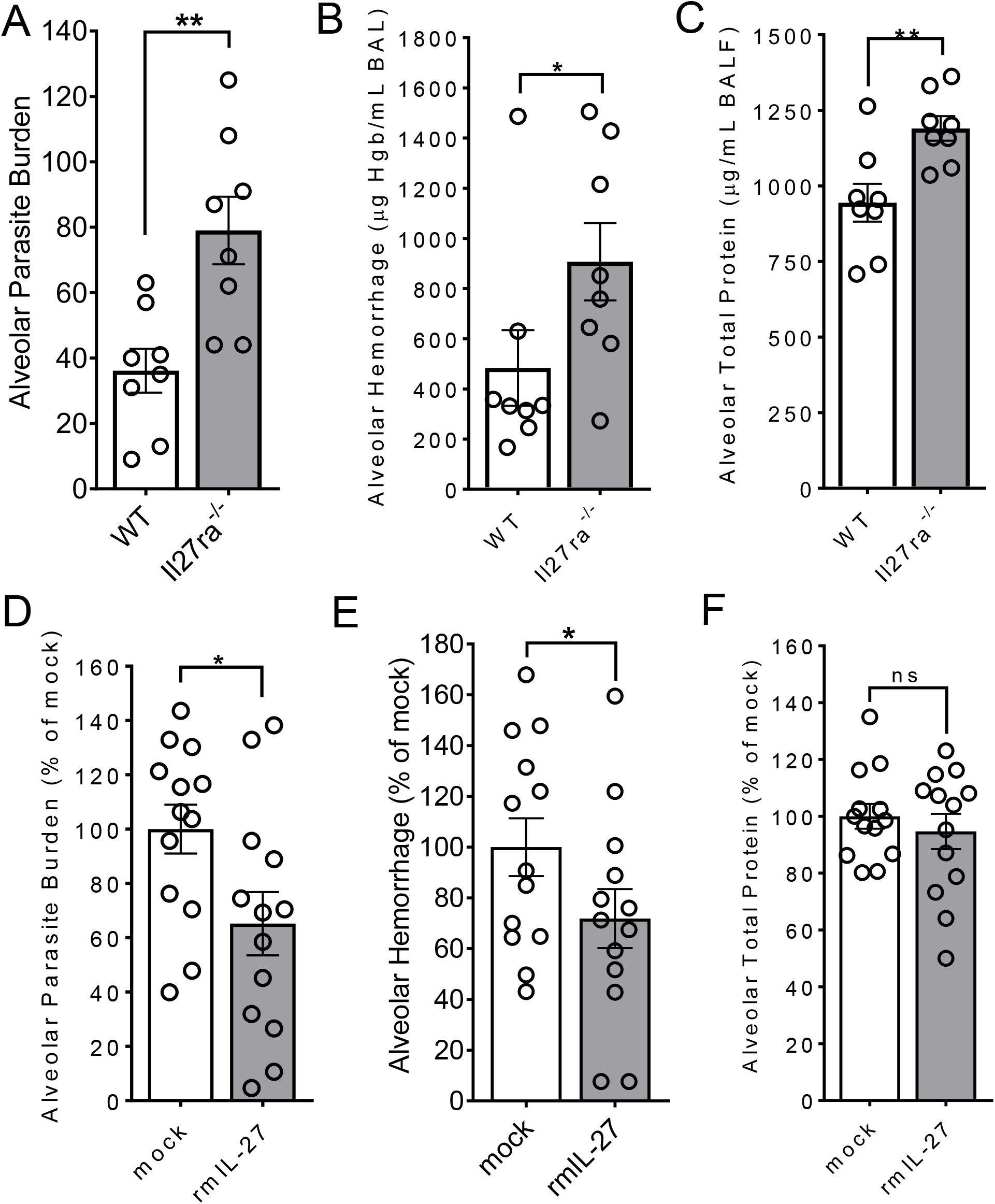
IL-27 enhances innate resistance to primary *N. brasiliensis* infection in the lungs. (**A-C**) *In vivo* comparison of susceptibility of Il27ra^−/−^ mice and wild type (WT; C57BL/6NJ) mice to primary *N. brasiliensis* infection (L3 n=500/mouse) in the lungs at 2 days after infection. (**A**) Alveolar parasite burden, (**B**) alveolar hemorrhage, and (**C**) alveolar total protein in BALF. (**D**-**F**) *In vivo* comparison of susceptibility of WT mice (C57BL/6J) administered with rmIL-27 (100 ng/mouse i.p., once daily on days 0-1) or mock control (0.1% BSA in PBS) during *N. brasiliensis* infection and analyzed after 2 days p.i. (**D**) Comparison of alveolar parasite burden, (**E**) alveolar hemorrhage, and (**F**) alveolar total protein in BALF. Data (A-F) are shown as mean ± SEM and each circle indicates an individual mouse, * *P*<0.05, ** *P*<0.01, ns: not significant.

Next, we tested whether administration of exogenous IL-27 would further reduce the severity of *N. brasiliensis* infection in the lungs of WT mice. Therefore, we administered 100 ng recombinant mouse IL-27 (rmIL-27) intraperitoneally on days 0 and 1 p.i. in C57BL/6J mice and compared alveolar parasite burden and injury with a carrier/mock alone group. The rmIL-27 treatment significantly reduced the alveolar parasite burden to 65% of mock controls (Fig. 4D). In addition, alveolar hemorrhage was significantly decreased to 72% of mock controls (Fig. 4E), while there was a slight, albeit statistically insignificant, decrease in total alveolar protein leakage (Fig. 4F). Taken together, these results indicate that both endogenous IL-27/IL-27RA signaling and therapeutic recombinant IL-27 enhance the innate resistance to primary *N. brasiliensis* infection during the early lung larval stage.

### IL-27 modulates the local presence of proinflammatory cytokines and chemokines

To characterize the influence of IL-27 on the local milieu of inflammatory mediators, we employed a high-sensitive, multiplexed bead-based assay to quantify the concentrations of 26 cytokines and chemokines (Fig. 5, S3, S4). Il27ra^−/−^ mice along with wild type (WT) control mice were inoculated with *N. brasiliensis* L3 and BALF was collected 2 days later. Significant differences were observed for 5 of 26 mediators. IL-6 concentrations were ~2-fold greater in Il27ra^−/−^ mice (Fig. 5A). TNFα was also significantly increased in Il27ra^−/−^ mice, although levels were near the lower detection limit of the assay, possibly owing to the time point chosen (Fig. 5A). The macrophage and T cell-driving chemokine, MCP-3, was 2-fold higher in Il27ra^−/−^ mice (Fig. 5A). In addition, we detected a trend towards lower concentrations for IL-23 (p = 0.09), a proinflammatory cytokine which is known to modulate T cell activity (Fig. 5A). Moreover, lower IL-10 concentrations and higher IL-17 amounts were detected (Fig. S3) in Il27ra^−/−^ mice, which is consistent with the known requirement of IL-27 signaling to induce IL-10 from T cells and to suppress IL-17 responses in other disease models (32–34).

**FIGURE 5.**
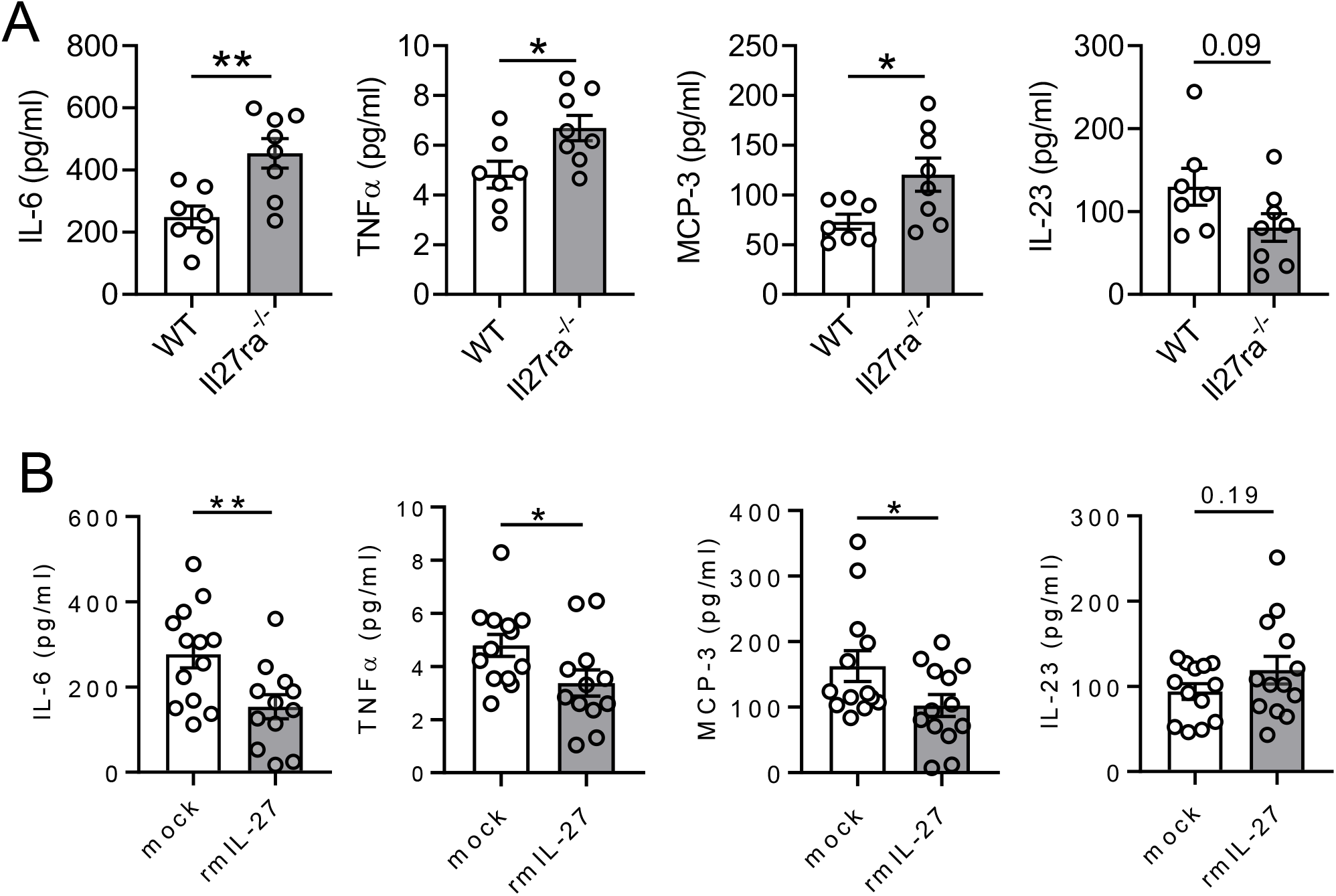
IL-27 regulates inflammatory mediators during primary pulmonary *N. brasiliensis* infection. (**A**) *In vivo* comparison of selected cytokines and chemokines of Il27ra^−/−^ mice and wild type control (C57BL/6NJ) mice to primary *N. brasiliensis* infection (L3 n=500/mouse) in the lungs (BALF) at 2 days after infection. (**B**) C57BL/6J mice administered with rmIL-27 (100 ng/mouse i.p., once daily on day 0 and day 1) or mock control (0.1% BSA in PBS) during *N. brasiliensis* infection and analyzed for the presence of inflammatory mediators after 2 days p.i in BALF. All data were obtained by multiplexed bead-based assay (Luminex-200). Graphs (A-B) are presented as mean ± SEM, were analyzed by two-tailed t-test and each circle indicates an individual mouse, * *P*<0.05, ** *P*<0.01.

To further elaborate on the findings with Il27ra^−/−^ mice, we measured inflammatory mediators in BALF of *N. brasiliensis* infected WT mice two days after a treatment regimen of rmIL-27 (100 ng i.p. on day 0 and day 1 p.i.) or mock injections. IL-6 was again observed to be present in abundant quantities, especially when considering the substantial dilution of alveolar lining fluid introduced by the lavage procedure (Fig. 5B). In contrast to the effect of IL-27 deficiency, IL-6 was suppressed by the addition of rmIL-27 (Fig. 5B). Likewise, mice that receive rmIL-27 were found to have lower TNFα and MCP-3 (Fig. 5B). Interestingly, no increase in IL-27 itself was detected in BALF 2 days after rmIL-27 injection (Fig. S4), suggesting clearance prior to that time point. In fact, we noticed a trend for suppressed (endogenous) IL-27 after injection, possibly attributable to a post excitation phenomenon.

### Absence of IL-27 signaling decreases the expansion of γδ T cells in the alveolar space during primary N. brasiliensis infection

As T cells by far have the highest expression if IL-27RA (Fig. 2C-E), and all three T cell subsets evaluated express IL-27RA (albeit highest on TCRβ^−^TCRγδ^+^ γδ T cells; Fig. 3), we next compared the percentage and number of all three T cells subsets in day 2 p.i. BAL between WT and Il27ra^−/−^ mice. Compared to WT mice we observed a significant 1.4-fold reduction in the percentage of TCRβ^−^TCRγδ^+^ γδ T cells, a 1.3-fold reduction in percentage of TCRβ^+^CD8^+^ CTLs and a slight increase in percentage of TCRβ^+^CD4^+^ Th cells (Fig. 6A, 6B) in Il27ra^−/−^ mice. Moreover, BALs from Il27ra^−/−^ mice exhibited a 1.5-fold reduction in the absolute number of TCRβ^−^TCRγδ^+^ γδ T cells, while the counts of TCRβ^+^CD4^+^ Th cells and TCRβ^+^CD8^+^ CTLs were unchanged. Thus, these findings support a role for IL-27 in enhancing the expansion of γδ T cells in the alveolar space during primary *N. brasiliensis* infections of the lungs.

**FIGURE 6.**
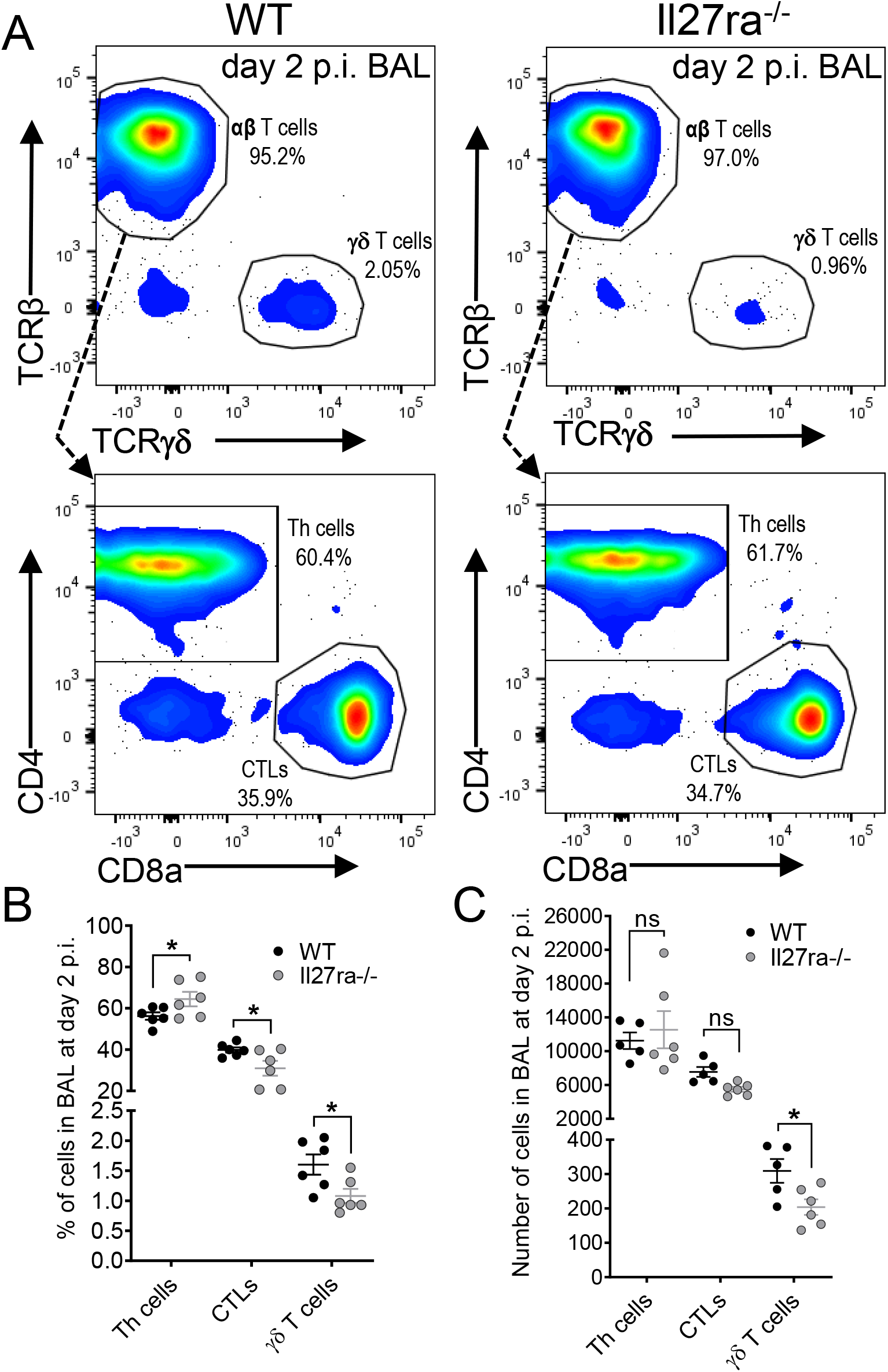
Il27ra^−/−^ mice have decreased presence of γδ T cells in the alveolar space during *N. brasiliensis* infection. (**A**) Representative flow cytometry plots of TCRβ^−^TCRγδ^+^ γδ T cells (top panel), and CD4^+^CD8^−^ Th cells and CD4^−^CD8^+^ CTLs (bottom panel), gated as in Fig. 3A, for WT (left panel) and Il27ra^−/−^ mice (right panel) in BAL on day 2 p.i. Percentages are indicated next to each gate. (**B**, **C**) Comparisons of frequencies (B) and absolute numbers (C) of CD4^+^CD8^−^ Th cells, CD4^−^CD8^+^ CTLs and TCRβ^−^TCRγδ^+^ γδ T cells in BAL of WT mice (n=5 mice/group; one mouse was removed due to extremely low infection, as determined by negligible hemorrhage) and Il27ra^−/−^ mice on day 2 p.i. (n=6 mice/group). Data (B, C) are shown as mean ± SEM, were analyzed by two-tailed t-test (WT vs. Il27ra^−/−^) and each symbol represents the value of an individual mouse, * *P*<0.05, ns: not significant.

## Discussion

In this report, we have identified a functional role for IL-27 signaling for protective immune defense against hookworm larvae, which migrate from the pulmonary vasculature into the airways. IL-27RA was expressed on all airway invading lymphocytes, albeit differentially based on cellular subset. IL-27RA was particularly abundant on γδ T cells compared to other innate and adaptive lymphocytes, suggesting this cell type as a prominent target for IL-27 mediated defense in the current setting. To this end, genetic deficiency of IL-27RA resulted in a greater parasite burden and more severe lung injury, whereas the opposite was true following IL-27 treatment. These findings were associated with differences in local proinflammatory cytokines and γδ T cell numbers in Il27ra^−/−^ mice.

IL-27-dependent changes occurred within the first two days of infection, suggesting that its beneficial roles stem from alterations in innate immunity, especially given the time required for antigen-specific T cell responses. Yet, cytokine-induced T cells can modulate inflammation independently of their T cell receptor (35, 36), such that their accumulation in the airspaces may indeed contribute to the inflammatory milieu in some capacity.

Innate immunity to helminths is widely accepted to be limited to expulsion, a canonical type 2 response orchestrated by Th2 cells, ILC2s and γδ T2 IECs that, in the case of hookworm infections, is not initiated until the adult stage of the life cycle in the small intestine (13, 16). Innate immune mechanisms responsible for defense against helminth infections within the lungs has received much less attention, likely due to the gut being the conserved final destination for the majority of helminths. We demonstrate that there are mechanisms of innate resistance to hookworm infection in the lungs that are not necessarily associated with type 2 responses. As a key booster of pulmonary resistance to hookworms, we found that IL-27 is transiently induced for expansion of γδ T cells in the alveolar space, where we anticipate their accumulation to directly and/or indirectly eradicate larvae prior to their transition to the small intestine. Our results strongly suggest that the role of IL-27 in the lung stage of hookworm infection is the opposite of the stage in the gut, being anti-parasitic for the former and pro-parasitic for the latter (22).

The downregulation of IL-6 production by IL-27 (Fig. 5) may be a direct effect, since lymphocytes can both produce and respond to IL-6 (37, 38). Alternatively, changes in IL-6 may occur as an indirect consequence of IL-27RA-expressing NK cells and T cells dispatching signals to control IL-6 synthesis in other non-lymphocytic cells. The precise role of IL-6 in lung injury and pulmonary inflammation appears to be somewhat dependent on the disease model (39, 40).

The chemokine MCP-3 (CCL7) shares 71% sequence similarity with MCP-1 (CCL2) and binds to the CCR2 receptor (41, 42). CCR2 is constitutively expressed not only on monocytes/macrophages, but also on T cells, including IL-17 producing γδ T cells (43). However, the higher concentrations of MCP-3 in Il27ra^−/−^ mice during hookworm infection seem to contradict the lower influx of γδ T cells in these mice (Fig. 5A vs. Fig. 6C). The increased MCP-3 may rather represent an ineffective compensatory feedback loop to bring back up T cell numbers, when too low. IL-27 itself is not a chemokine, but gp130-induced JAK/STAT1/STAT3 signaling may regulate expression of chemokines, chemokine receptors and T cell migration (44, 45). Another possibility is that elevated MCP-3 is secondary to the increase in pathogen burden resulting from IL-27 deficiency, making the precise roles of MCP-3 speculative at present.

IL-27 is almost exclusively produced by macrophages and dendritic cells (20, 46). Here, we have detected IL-27RA expression on all lung lymphocyte subpopulations during hookworm infection, which is consistent with abundant evidence highlighting the responsiveness and functional roles of IL-27 in T cells and B cells (20, 47). On the other hand, we did not observe IL-27RA expression on mouse myeloid cells in lungs. In fact, the expression of IL-27RA on myeloid cells appears to occur in dependency of cell maturation and species (48–52). IL-27RA expression and IL-27 responsiveness exist for human neutrophils and human monocytes, whereas mouse macrophages only display minimal responsiveness (53, 54). We caution that none of the earlier reports have assessed IL-27RA expression in a cell-specific capacity (as accomplished by flow cytometry in the present study), but instead relied on RT-PCR or western blotting of cell lysates, making conclusive determination of cellular source somewhat speculative.

The regulatory role of IL-27 for the host immune defense against helminths does not appear to be limited to hookworms. Dual deficiency of IL-27RA and IL-10 rescues the great susceptibility of IL-10 single knockout mice for intestinal pathology and infection caused by the nematode *Trichuris muris* (55). *Strongyloides stercolaris,* the causative threadworm of Strongyloidiasis, infects more than 50 million people worldwide and infection results in a moderate but significant increase of IL-27 in human plasma, which decreases after anthelmintic treatment (56).

In human whole blood cultures of infected individuals re-stimulated with recombinant *S. stercolaris* NIE antigen, the neutralization of IL-27 using antibodies increased the frequencies of all CD4^+^ T helper cell subsets (Th1-Th22), CD8^+^ T cells and modulated cytokine levels (57). *Ascaris lubricoides* antigen induced IL-27 release from PBMCs in adults and the elderly as compared to neonates and children of an endemic cohort from Sub-Saharan Africa, with a 30% prevalence of hook worm infections and multiple parasite infections (58). Altogether, these observations suggest an important influence of IL-27 on host outcome in a variety of settings of helminthic infections.

The mechanisms of innate immune recognition for helminths remain unclear. No dedicated class of pattern recognition receptors has been identified so far. Helminth-derived chitin, proteoglycans, lipids and excretory-secretory products may be recognized by TLRs and C-type lectins (59, 60). In addition, danger associated molecular patterns (DAMPs) when released during helminth-induced tissue injury could act as endogenous ligands for TLRs (and other receptors), thereby inducing MYD88/TRIF-dependent IL-27 production (46, 61, 62). Furthermore, γδ T cells can be activated by DAMPs from mitochondria (63).

While we describe in this report that recombinant IL-27 reinforced the anthelmintic host defense in the lungs, the feasibility of proposing its administration as an effective therapy is highly speculative. First, the efficacy observed in our studies was less than 50% for reduction of the lung larvae burden in IL-27-treated mice, although this could be improved by further dose optimization. Secondly, the skin penetration of hookworm larvae in humans usually is unnoticed, such that any window of therapeutic efficacy may be too difficult to rely on. Thirdly, another report showed that a non-viral minicircle DNA vector injected intra-venously for recombinant expression of IL-27 resulted in higher numbers of adult *N. brasiliensis* in the intestinal tract (22). This implies that the role of IL-27 could be either protective or detrimental depending on the stage of the hookworm infection cycle and host tissue environment. It appears that more sophisticated manipulations would be needed to specifically enhance IL-27-dependent protective immunity in the lung. IL-27 may be an interesting factor to explore for developing adjuvants for vaccines that target the early lung stage of human hookworms (11). Vaccine adjuvants would need to be tailored to locally induce IL-27 (27–29), or to up-regulate IL-27RA expression and IL-27 responsiveness of lung resident lymphocyte subpopulations (64).

In conclusion, the presented work expands on the emerging role of IL-27 as a critical factor of host defense and immune regulation against hookworm infections. The pulmonary stage of the parasitic infection cycle remains an understudied area and we highlight the importance to better understand the lung lymphocyte-specific host response. In the future, more research will be needed to fully uncover the intricate molecular mechanisms of host-helminth interactions as a basis for developing innovative treatment approaches against this neglected tropical disease.

## Acknowledgments

We thank Dr. Joseph Urban Jr. (USDA, Beltsville, MD) for kindly providing the murine hookworm *N. brasiliensis*. This work was supported by the National Institutes of Health (1R01HL141513, 1R01HL139641 to M.B.), the Federal Ministry of Education and Research (01EO1503 to M.B. and C.R.), the Deutsche Forschungsgemeinschaft (BO3482/3-3, BO3482/4-1 to M.B. and RE-3450/5-2 to C.R.), a Marie Curie Career Integration Grant of the European Union (Project 334486 to M.B.), a Clinical Research Fellowship of the European Hematology Association (to M.B.) and a project grant from the Boehringer Ingelheim Foundation (Consortium Grant “Novel and neglected cardiovascular risk factors” to C.R.). C.R. was awarded a fellowship from the Gutenberg Research College. The authors are responsible for the contents of this publication.

## Author Contributions

J.B.N. and A.S. designed and performed experiments and analyzed data. J.P. and C.R. contributed to experimental designs and provided helpful comments. M.B. conceived and supervised the study, designed experiments, interpreted data and provided funding. J.B.N. and M.B. wrote the manuscript, which was further edited by A.S., L.J.Q. and C.R.

## Correspondence and requests for materials

Requests for materials and correspondence should be addressed to M.B.

---

**FIGURE S1.**
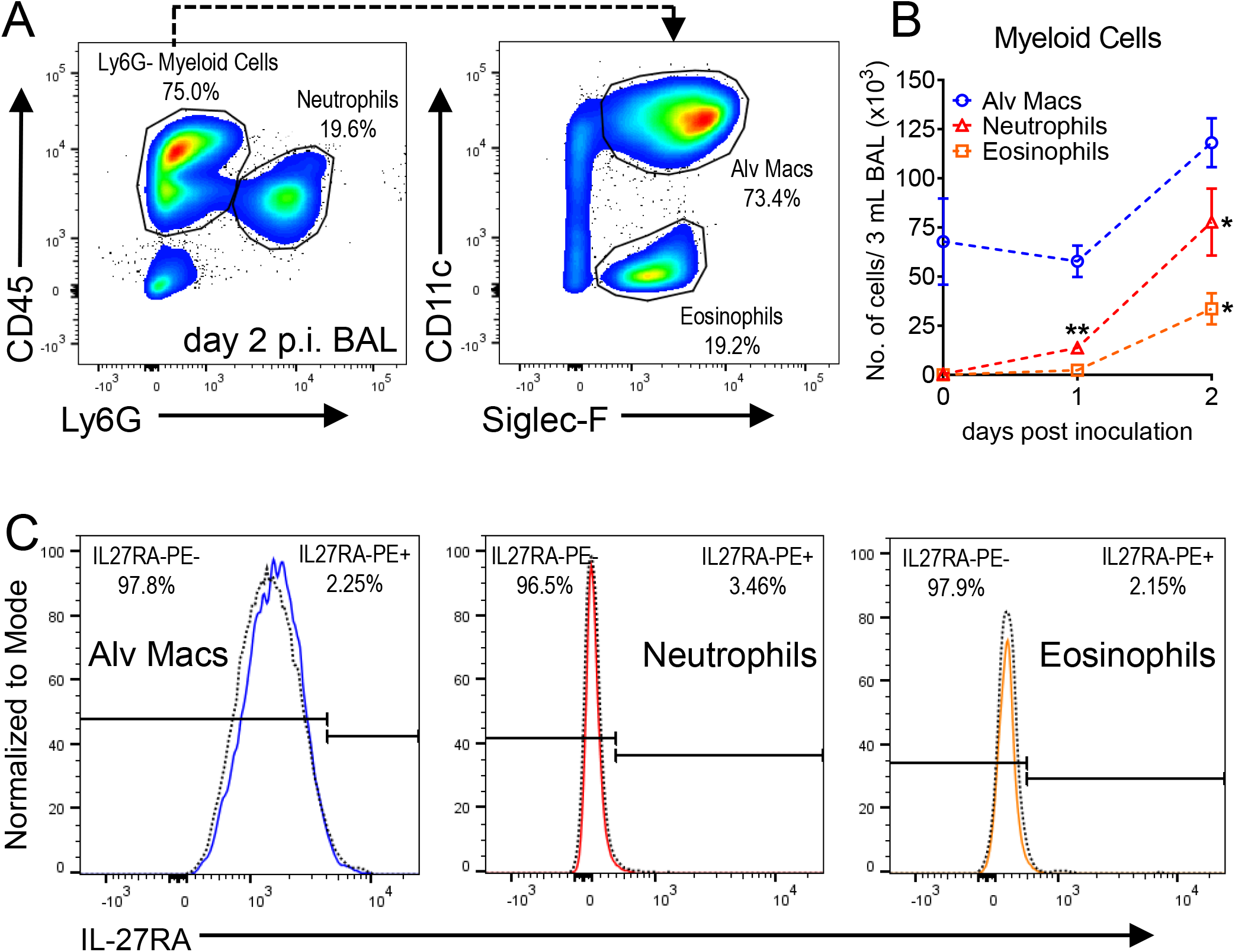
IL-27RA is not expressed on alveolar macrophages, neutrophils or eosinophils from the alveolar space during *N. brasiliensis* infection. (**A**) Flow cytometry plots including gating strategy of CD45 versus Ly6G pre-gated on single myeloid cells (left) and CD11c versus Siglec-F on Ly6G^−^ myeloid cells (right). Percentages are indicated next to each gate. (**B**) Numbers of Ly6G^+^ neutrophils, Ly6G^−^CD11c^+^Siglec-F^+^ alveolar macrophages and Ly6G^−^CD11c^−^Siglec-F^+^ eosinophils on day 0 (PBS-inoculated/uninfected), day 1 and day 2 post-inoculation (n=3 mice/group) in BAL; day 0 vs. day 1 and day 0 vs. day 2 using two-tailed t-test, data shows mean ± SEM. (**C**) Histograms of IL-27RA expression on Ly6G^−^CD11c^+^Siglec-F^+^ alveolar macrophages (left), Ly6G^+^ neutrophils (middle) and Ly6G^−^CD11c^−^ Siglec-F^+^ eosinophils (right) on day 2 p.i. The dotted line indicates isotype-FMO control. The percentages of IL-27RA^−^ and IL-27RA^+^ cells are indicated in the upper left and right corners, respectively.

**FIGURE S2.**
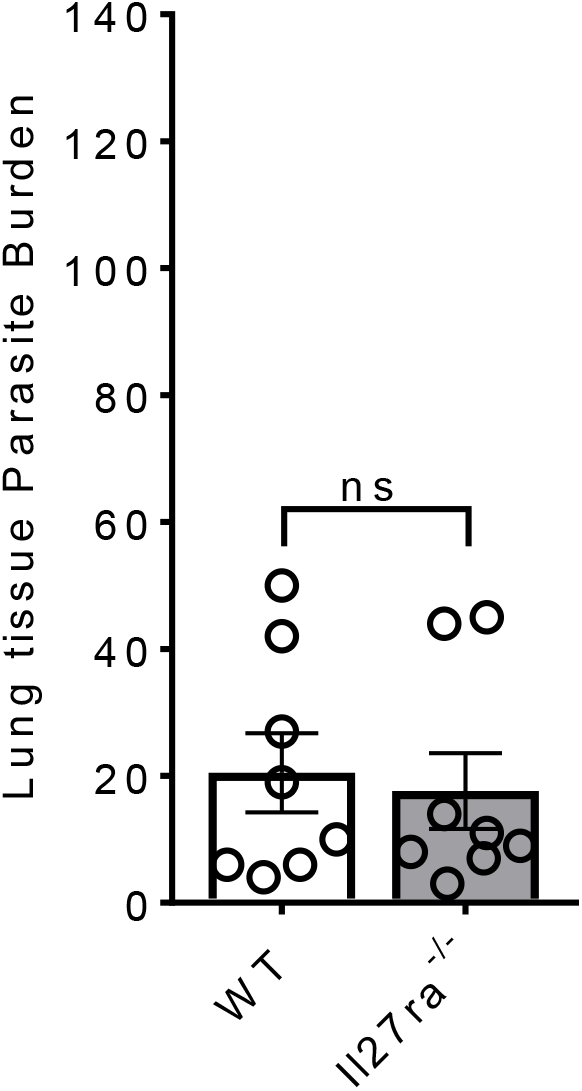
Parasite burdens are similar between wild type mice compared to Il27ra^−/−^ mice in the lung tissues during *N. brasiliensis* infection. Wild type mice (C57BL/6NJ) and Il27ra^−/−^ mice were inoculated s.c. with L3 larvae (n=500/mouse). Parasite numbers were counted in whole lung tissues after removal of intra-airway parasites by multiple BAL on day 2 of infection. Lungs were minced from the same experiments as shown in Fig. 4A. Comparison of mean ± SEM analyzed by two-tailed t-test and each circle indicates the value of an individual mouse, ns: not significant.

**FIGURE S3.**
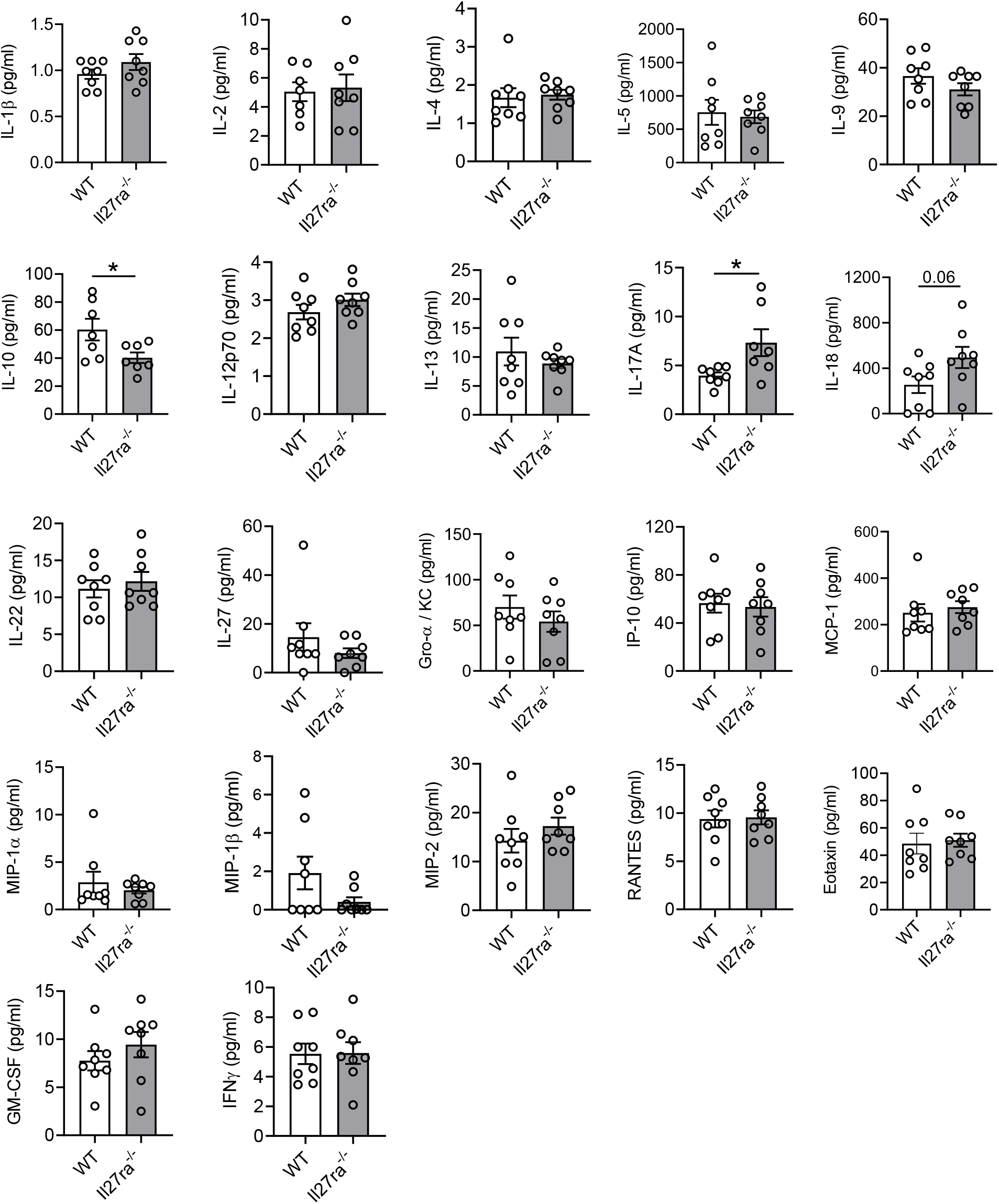
Cytokines and chemokines in brochoalveolar lavage fluids of Il27ra^−/−^ mice. *In vivo* comparison of different cytokines and chemokines of Il27ra^−/−^ mice and wild type (WT; C57BL/6NJ) mice during primary *N. brasiliensis* infection (L3 n=500/mouse) in BALF after 2 days, bead-based multiplex assay. Data are shown as mean ± SEM and each circle represents an individual mouse, two-tailed t-test, * *P*<0.05, ** *P*<0.01.

**FIGURE S4.**
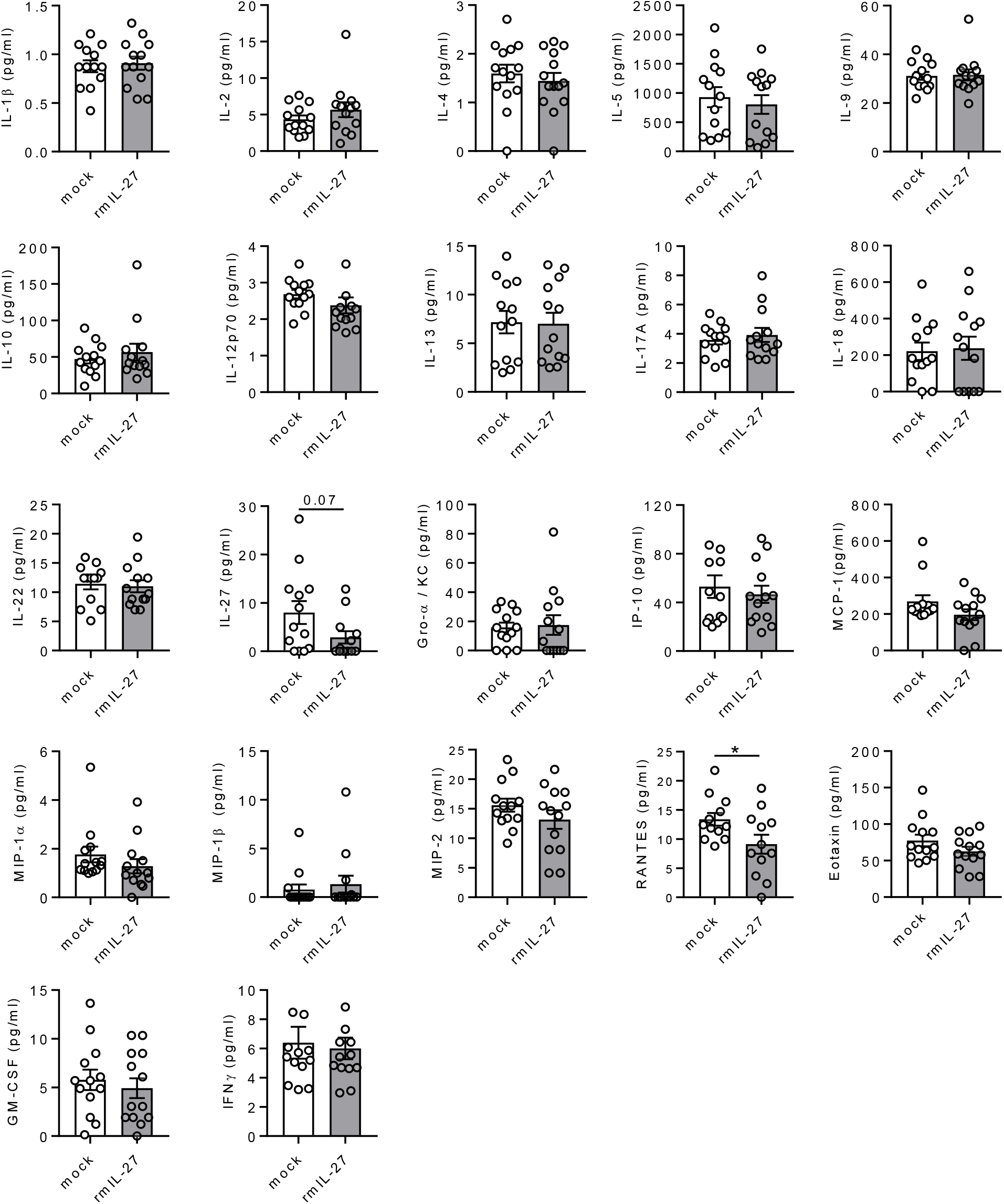
Cytokines and chemokines in brochoalveolar lavage fluids after treatment with recombinant mouse IL-27. *In vivo* comparison of mediators in BALF from WT mice (C57BL/6J) administered with rmIL-27 (100 ng/mouse i.p.) or mock control (0.1% BSA in PBS) on day 0 and day 1 of *N. brasiliensis* infection and analyzed after 2 days p.i. Data are shown as mean ± SEM and each circle indicates an individual mouse, two-tailed t-test, * *P*<0.05.

## References

1. Loukas, A., J. Hotez, D. Diemert, M. Yazdanbakhsh, J. S. McCarthy, R. Correa-Oliveira, J. Croese, and J. M. Bethony. 2016. Hookworm infection. Nat Rev Dis Primers 2: 16088.

2. Pullan, R. L., J. L. Smith, R. Jasrasaria, and S. J. Brooker. 2014. Global numbers of infection and disease burden of soil transmitted helminth infections in 2010. Parasit Vectors 7: 37.

3. Herricks, J. R., P. J. Hotez, V. Wanga, L. E. Coffeng, J. A. Haagsma, M. G. Basanez, G. Buckle, C. M. Budke, H. Carabin, E. M. Fevre, T. Furst, Y. A. Halasa, C. H. King, M. E. Murdoch, K. D. Ramaiah, D. S. Shepard, W. A. Stolk, E. A. Undurraga, J. D. Stanaway, M. Naghavi, and C. J. L. Murray. 2017. The global burden of disease study 2013: What does it mean for the NTDs? PLoS Negl Trop Dis 11: e0005424.

4. Brooker, S., P. J. Hotez, and D. A. Bundy. 2008. Hookworm-related anaemia among pregnant women: a systematic review. PLoSNegl Trop Dis 2: e291.

5. Smith, J. L., and S. Brooker. 2010. Impact of hookworm infection and deworming on anaemia in non-pregnant populations: a systematic review. Trop Med Int Health 15: 776–795.

6. Hotez, P. J., S. Brooker, J. M. Bethony, M. E. Bottazzi, A. Loukas, and S. Xiao. 2004. Hookworm infection. N Engl J Med 351: 799–807.

7. Kassebaum, N. J., and G. B. D. A. Collaborators. 2016. The Global Burden of Anemia. Hematol Oncol Clin North Am 30: 247–308.

8. Hotez, P. J., D. A. P. Bundy, K. Beegle, S. Brooker, L. Drake, N. de Silva, A. Montresor, D. Engels, M. Jukes, L. Chitsulo, J. Chow, R. Laxminarayan, C. Michaud, J. Bethony, R. Correa-Oliveira, X. Shuhua, A. Fenwick, and L. Savioli. 2006. Helminth Infections: Soil-transmitted Helminth Infections and Schistosomiasis. In Disease Control Priorities in Developing Countries. nd, D. T. Jamison, J. G. Breman, A. R. Measham, G. Alleyne, M. Claeson, D. B. Evans, P. Jha, A. Mills, and P. Musgrove, eds, Washington (DC).

9. Anderson, R., J. Truscott, and T. D. Hollingsworth. 2014. The coverage and frequency of mass drug administration required to eliminate persistent transmission of soil-transmitted helminths. Philos Trans R Soc Lond B Biol Sci 369: 20130435.

10. Bethony, J., S. Brooker, M. Albonico, S. M. Geiger, A. Loukas, D. Diemert, and P. J. Hotez. 2006. Soil-transmitted helminth infections: ascariasis, trichuriasis, and hookworm. Lancet 367: 1521–1532.

11. Noon, J. B., and R. V. Aroian. 2017. Recombinant subunit vaccines for soil-transmitted helminths. Parasitology 144: 1845–1870.

12. Camberis, M., G. Le Gros, and J. Urban, Jr. 2003. Animal model of Nippostrongylus brasiliensis and Heligmosomoides polygyrus. Curr Protoc Immunol Chapter 19: Unit 19 12.

13. Nair, M. G., and D. R. Herbert. 2016. Immune polarization by hookworms: taking cues from T helper type 2, type 2 innate lymphoid cells and alternatively activated macrophages. Immunology 148: 115–124.

14. Sutherland, T. E., N. Logan, D. Ruckerl, A. A. Humbles, S. M. Allan, V. Papayannopoulos, B. Stockinger, R. M. Maizels, and J. E. Allen. 2014. Chitinase-like proteins promote IL-17-mediated neutrophilia in a tradeoff between nematode killing and host damage. Nat Immunol 15: 1116–1125.

15. Cardoso, V., J. Chesne, H. Ribeiro, B. Garcia-Cassani, T. Carvalho, T. Bouchery, K. Shah, N. L. Barbosa-Morais, N. Harris, and H. Veiga-Fernandes. 2017. Neuronal regulation of type 2 innate lymphoid cells via neuromedin U. Nature 549: 277–281.

16. Inagaki-Ohara, K., Y. Sakamoto, T. Dohi, and A. L. Smith. 2011. gammadelta T cells play a protective role during infection with Nippostrongylus brasiliensis by promoting goblet cell function in the small intestine. Immunology 134: 448–458.

17. Pflanz, S., J. C. Timans, J. Cheung, R. Rosales, H. Kanzler, J. Gilbert, L. Hibbert, T. Churakova, M. Travis, E. Vaisberg, W. M. Blumenschein, J. D. Mattson, J. L. Wagner, W. To, S. Zurawski, T. K. McClanahan, D. M. Gorman, J. F. Bazan, R. de Waal Malefyt, D. Rennick, and R. A. Kastelein. 2002. IL-27, a heterodimeric cytokine composed of EBI3 and p28 protein, induces proliferation of naive CD4(+) T cells. Immunity 16: 779–790.

18. Collison, L. W., C. J. Workman, T. T. Kuo, K. Boyd, Y. Wang, K. M. Vignali, R. Cross, D. Sehy, R. S. Blumberg, and D. A. Vignali. 2007. The inhibitory cytokine IL-35 contributes to regulatory T-cell function. Nature 450: 566–569.

19. Hamano, S., K. Himeno, Y. Miyazaki, K. Ishii, A. Yamanaka, A. Takeda, M. Zhang, H. Hisaeda, T. W. Mak, A. Yoshimura, and H. Yoshida. 2003. WSX-1 is required for resistance to Trypanosoma cruzi infection by regulation of proinflammatory cytokine production. Immunity 19: 657–667.

20. Bosmann, M., and P. A. Ward. 2013. Modulation of inflammation by interleukin-27. J Leukoc Biol 94: 1159–1165.

21. Hunter, C. A., and R. Kastelein. 2012. Interleukin-27: balancing protective and pathological immunity. Immunity 37: 960–969.

22. McHedlidze, T., M. Kindermann, A. T. Neves, D. Voehringer, M. F. Neurath, and S. Wirtz. 2016. IL-27 suppresses type 2 immune responses in vivo via direct effects on group 2 innate lymphoid cells. Mucosal Immunol 9: 1384–1394.

23. Artis, D., A. Villarino, M. Silverman, W. He, E. M. Thornton, S. Mu, S. Summer, T. M. Covey, E. Huang, H. Yoshida, G. Koretzky, M. Goldschmidt, G. D. Wu, F. de Sauvage, H. R. Miller, C. J. Saris, P. Scott, and C. A. Hunter. 2004. The IL-27 receptor (WSX-1) is an inhibitor of innate and adaptive elements of type 2 immunity. J Immunol 173: 5626–5634.

24. Filbey, K., T. Bouchery, and G. Le Gros. 2018. The role of ILC2 in hookworm infection. Parasite Immunol 40.

25. Fallon, P. G., S. J. Ballantyne, N. E. Mangan, J. L. Barlow, A. Dasvarma, D. R. Hewett, A. McIlgorm, H. E. Jolin, and A. N. McKenzie. 2006. Identification of an interleukin (IL)-25-dependent cell population that provides IL-4, IL-5, and IL-13 at the onset of helminth expulsion. J Exp Med 203: 1105–1116.

26. Moro, K., H. Kabata, M. Tanabe, S. Koga, N. Takeno, M. Mochizuki, K. Fukunaga, K. Asano, T. Betsuyaku, and S. Koyasu. 2016. Interferon and IL-27 antagonize the function of group 2 innate lymphoid cells and type 2 innate immune responses. Nat Immunol 17: 76–86.

27. Pennock, N. D., L. Gapin, and R. M. Kedl. 2014. IL-27 is required for shaping the magnitude, affinity distribution, and memory of T cells responding to subunit immunization. Proc Natl Acad Sci U S A 111: 16472–16477.

28. Kilgore, A. M., N. D. Pennock, and R. M. Kedl. 2020. cDC1 IL-27p28 Production Predicts Vaccine-Elicited CD8(+) T Cell Memory and Protective Immunity. J Immunol 204: 510–517.

29. Coffman, R. L., A. Sher, and R. A. Seder. 2010. Vaccine adjuvants: putting innate immunity to work. Immunity 33: 492–503.

30. Oshiro, I., T. Takenaka, and J. Maeda. 1982. New method for hemoglobin determination by using sodium lauryl sulfate (SLS). Clin Biochem 15: 83–88.

31. Bosmann, M., N. F. Russkamp, V. R. Patel, F. S. Zetoune, J. V. Sarma, and P. A. Ward. 2011. The outcome of polymicrobial sepsis is independent of T and B cells. Shock 36: 396–401.

32. Batten, M., J. Li, S. Yi, N. M. Kljavin, D. M. Danilenko, S. Lucas, J. Lee, F. J. de Sauvage, and N. Ghilardi. 2006. Interleukin 27 limits autoimmune encephalomyelitis by suppressing the development of interleukin 17-producing T cells. Nature Immunology 7: 929–936.

33. Stumhofer, J. S., A. Laurence, E. H. Wilson, E. Huang, C. M. Tato, L. M. Johnson, A. V. Villarino, Q. Huang, A. Yoshimura, D. Sehy, C. J. Saris, J. J. O’Shea, L. Hennighausen, M. Ernst, and C. A. Hunter. 2006. Interleukin 27 negatively regulates the development of interleukin 17-producing T helper cells during chronic inflammation of the central nervous system. Nat Immunol 7: 937–945.

34. Stumhofer, J. S., J. S. Silver, A. Laurence, P. M. Porrett, T. H. Harris, L. A. Turka, M. Ernst, C. J. Saris, J. J. O’Shea, and C. A. Hunter. 2007. Interleukins 27 and 6 induce STAT3-mediated T cell production of interleukin 10. Nat Immunol 8: 1363–1371.

35. Schmidt-Wolf, I. G., R. S. Negrin, H. P. Kiem, K. G. Blume, and I. L. Weissman. 1991. Use of a SCID mouse/human lymphoma model to evaluate cytokine-induced killer cells with potent antitumor cell activity. J Exp Med 174: 139–149.

36. Meresse, B., Z. Chen, C. Ciszewski, M. Tretiakova, G. Bhagat, T. N. Krausz, D. H. Raulet, L. L. Lanier, V. Groh, T. Spies, E. C. Ebert, P. H. Green, and B. Jabri. 2004. Coordinated induction by IL15 of a TCR-independent NKG2D signaling pathway converts CTL into lymphokine-activated killer cells in celiac disease. Immunity 21: 357–366.

37. Avanzi, G. C., A. Cessano, M. F. Brizzi, S. C. Clark, L. Pegoraro, and L. Matera. 1989. Biological and molecular evidence for the production of IL-6 by human natural killer cells in culture. Life Sci 45: 2621–2626.

38. Kuhn, K. A., N. A. Manieri, T. C. Liu, and T. S. Stappenbeck. 2014. IL-6 stimulates intestinal epithelial proliferation and repair after injury. PLoS One 9: e114195.

39. Saito, F., S. Tasaka, K. Inoue, K. Miyamoto, Y. Nakano, Y. Ogawa, W. Yamada, Y. Shiraishi, N. Hasegawa, S. Fujishima, H. Takano, and A. Ishizaka. 2008. Role of interleukin-6 in bleomycin-induced lung inflammatory changes in mice. Am J Respir Cell Mol Biol 38: 566–571.

40. Bhargava, R., W. Janssen, C. Altmann, A. Andres-Hernando, K. Okamura, R. W. Vandivier, N. Ahuja, and S. Faubel. 2013. Intratracheal IL-6 protects against lung inflammation in direct, but not indirect, causes of acute lung injury in mice. PLoS One 8: e61405.

41. Sozzani, S., D. Zhou, M. Locati, M. Rieppi, P. Proost, M. Magazin, N. Vita, J. van Damme, and A. Mantovani. 1994. Receptors and transduction pathways for monocyte chemotactic protein-2 and monocyte chemotactic protein-3. Similarities and differences with MCP-1. J Immunol 152: 3615–3622.

42. Franci, C., L. M. Wong, J. Van Damme, P. Proost, and I. F. Charo. 1995. Monocyte chemoattractant protein-3, but not monocyte chemoattractant protein-2, is a functional ligand of the human monocyte chemoattractant protein-1 receptor. J Immunol 154: 6511–6517.

43. McKenzie, D. R., E. E. Kara, C. R. Bastow, T. S. Tyllis, K. A. Fenix, C. E. Gregor, J. J. Wilson, R. Babb, J. C. Paton, A. Kallies, S. L. Nutt, A. Brustle, M. Mack, I. Comerford, and S. R. McColl. 2017. IL-17-producing gammadelta T cells switch migratory patterns between resting and activated states. Nat Commun 8: 15632.

44. McLoughlin, R. M., B. J. Jenkins, D. Grail, A. S. Williams, C. A. Fielding, C. R. Parker, M. Ernst, N. Topley, and S. A. Jones. 2005. IL-6 trans-signaling via STAT3 directs T cell infiltration in acute inflammation. Proc Natl Acad Sci U S A 102: 9589–9594.

45. Barbi, J., S. Oghumu, C. M. Lezama-Davila, and A. R. Satoskar. 2007. IFN-gamma and STAT1 are required for efficient induction of CXC chemokine receptor 3 (CXCR3) on CD4+ but not CD8+ T cells. Blood 110: 2215–2216.

46. Bosmann, M., N. F. Russkamp, B. Strobl, J. Roewe, L. Balouzian, F. Pache, M. P. Radsak, N. van Rooijen, F. S. Zetoune, J. V. Sarma, G. Nunez, M. Muller, P. J. Murray, and P. A. Ward. 2014. Interruption of macrophage-derived IL-27(p28) production by IL-10 during sepsis requires STAT3 but not SOCS3. J Immunol 193: 5668–5677.

47. Vijayan, D., N. Mohd Redzwan, D. T. Avery, R. C. Wirasinha, R. Brink, G. Walters, S. Adelstein, M. Kobayashi, P. Gray, M. Elliott, M. Wong, C. King, C. G. Vinuesa, N. Ghilardi, C. S. Ma, S. G. Tangye, and M. Batten. 2016. IL-27 Directly Enhances Germinal Center B Cell Activity and Potentiates Lupus in Sanroque Mice. J Immunol 197: 3008–3017.

48. Pflanz, S., L. Hibbert, J. Mattson, R. Rosales, E. Vaisberg, J. F. Bazan, J. H. Phillips, T. K. McClanahan, R. de Waal Malefyt, and R. A. Kastelein. 2004. WSX-1 and glycoprotein 130 constitute a signal-transducing receptor for IL-27. J Immunol 172: 2225–2231.

49. Kalliolias, G. D., and L. B. Ivashkiv. 2008. IL-27 activates human monocytes via STAT1 and suppresses IL-10 production but the inflammatory functions of IL-27 are abrogated by TLRs and p38. J Immunol 180: 6325–6333.

50. Holscher, C., A. Holscher, D. Ruckerl, T. Yoshimoto, H. Yoshida, T. Mak, C. Saris, and S. Ehlers. 2005. The IL-27 receptor chain WSX-1 differentially regulates antibacterial immunity and survival during experimental tuberculosis. J Immunol 174: 3534–3544.

51. Zhao, X., S. M. Ting, C. H. Liu, G. Sun, M. Kruzel, M. Roy-O’Reilly, and J. Aronowski. 2017. Neutrophil polarization by IL-27 as a therapeutic target for intracerebral hemorrhage. Nat Commun 8: 602.

52. Seita, J., M. Asakawa, J. Ooehara, S. Takayanagi, Y. Morita, N. Watanabe, K. Fujita, M. Kudo, J. Mizuguchi, H. Ema, H. Nakauchi, and T. Yoshimoto. 2008. Interleukin-27 directly induces differentiation in hematopoietic stem cells. Blood 111: 1903–1912.

53. Li, J. P., H. Wu, W. Xing, S. G. Yang, S. H. Lu, W. T. Du, J. X. Yu, F. Chen, L. Zhang, and Z. C. Han. 2010. Interleukin-27 as a negative regulator of human neutrophil function. Scand J Immunol 72: 284–292.

54. Ruckerl, D., M. Hessmann, T. Yoshimoto, S. Ehlers, and C. Holscher. 2006. Alternatively activated macrophages express the IL-27 receptor alpha chain WSX-1. Immunobiology 211: 427–436.

55. Villarino, A. V., D. Artis, J. S. Bezbradica, O. Miller, C. J. Saris, S. Joyce, and C. A. Hunter. 2008. IL-27R deficiency delays the onset of colitis and protects from helminth-induced pathology in a model of chronic IBD. Int Immunol 20: 739–752.

56. Anuradha, R., S. Munisankar, Y. Bhootra, J. Jagannathan, C. Dolla, P. Kumaran, K. Shen, T. B. Nutman, and S. Babu. 2016. Systemic Cytokine Profiles in Strongyloides stercoralis Infection and Alterations following Treatment. Infect Immun 84: 425–431.

57. Anuradha, R., S. Munisankar, Y. Bhootra, C. Dolla, P. Kumaran, T. B. Nutman, and S. Babu. 2017. Modulation of CD4(+) and CD8(+) T Cell Function and Cytokine Responses in Strongyloides stercoralis Infection by Interleukin-27 (IL-27) and IL-37. Infect Immun 85.

58. Hegewald, J., R. G. Gantin, C. J. Lechner, X. Huang, A. Agosssou, Y. F. Agbeko, P. T. Soboslay, and C. Kohler. 2015. Cellular cytokine and chemokine responses to parasite antigens and fungus and mite allergens in children co-infected with helminthes and protozoa parasites. JInflamm (Lond) 12: 5.

59. Perrigoue, J. G., F. A. Marshall, and D. Artis. 2008. On the hunt for helminths: innate immune cells in the recognition and response to helminth parasites. Cell Microbiol 10: 1757–1764.

60. McGuinness, D. H., P. K. Dehal, and R. J. Pleass. 2003. Pattern recognition molecules and innate immunity to parasites. Trends Parasitol 19: 312–319.

61. Bosmann, M., M. D. Haggadone, M. R. Hemmila, F. S. Zetoune, J. V. Sarma, and P. A. Ward. 2012. Complement activation product C5a is a selective suppressor of TLR4-induced, but not TLR3-induced, production of IL-27(p28) from macrophages. J Immunol 188: 5086–5093.

62. Bosmann, M., B. Strobl, N. Kichler, D. Rigler, J. J. Grailer, F. Pache, P. J. Murray, M. Muller, and P. A. Ward. 2014. Tyrosine kinase 2 promotes sepsis-associated lethality by facilitating production of interleukin-27. J Leukoc Biol 96: 123–131.

63. Schwacha, M. G., M. Rani, S. E. Nicholson, A. M. Lewis, T. L. Holloway, S. Sordo, and A. P. Cap. 2016. Dermal gammadelta T-Cells Can Be Activated by Mitochondrial Damage-Associated Molecular Patterns. PLoS One 11: e0158993.

64. Szabo, P. A., M. Miron, and D. L. Farber. 2019. Location, location, location: Tissue resident memory T cells in mice and humans. Sci Immunol 4

